# Single-cell RNA-sequencing of cerebral spinal fluid identifies circulating tumour cells in children with brain cancer

**DOI:** 10.64898/2025.12.18.694991

**Authors:** Chelsea Mayoh, Wenyan Li, Mojgan Toumari, Sajad Razavi Bazaz, Anne-Lise Gerard, Aileen Lowe, Paulette Barahona, Loretta MS Lau, Jordan Staunton, Erica Jacobson, Sumanth Nagabushan, Neevika Manoharan, Ruth Mitchell, Faustine Ong, Marie Wong-Erasmus, Paul G Ekert, Vanessa Tyrrell, Michelle Haber, David S Ziegler, Mark J Cowley, Marion K Mateos, Robert Salomon

**Affiliations:** Children’s Cancer Institute, Lowy Cancer Research Centre, UNSW, Kensington, NSW, Australia; School of Clinical Medicine, UNSW Medicine & Health, UNSW Sydney, Kensington, NSW, Australia; Kids Cancer Centre, Sydney Children’s Hospital, Randwick, NSW, Australia; Cancer Centre for Children, Children’s Hospital at Westmead, Westmead, NSW, Australia; Murdoch Children’s Research Institute, Royal Children’s Hospital, Parkville, Victoria, Australia; Cancer Immunology Program, Peter MacCallum Cancer Centre, Parkville, Victoria, Australia; The Sir Peter MacCallum Department of Oncology, University of Melbourne, Parkville, Victoria, Australia

## Abstract

Paediatric central nervous system (CNS) tumours are the leading cause of cancer-related death in children, yet disease monitoring remains challenging. Conventional approaches, including imaging and cytology, lack sensitivity, delaying intervention. Liquid biopsy offers a minimally invasive alternative, but the utility of circulating tumour cells (CTCs) in paediatric CNS tumours as biomarkers is poorly defined. We developed a CTC detection and characterisation workflow from cerebrospinal fluid (CSF) utilising single-cell RNA-sequencing (scRNA-seq) and applied this to ten CNS tumour subtypes in 16 patients. CTCs were identified in all cases, with higher burdens in pineoblastoma, medulloblastoma and atypical teratoid rhabdoid tumours. Longitudinal profiling revealed CTC dynamics correlated with clinical disease course and anticipated relapse. Critically, scRNA-seq uncovered a sub-clonal canonical driver alteration at diagnosis that only became detectable by bulk RNA-seq at progression, underscoring its potential to resolve clonal dynamics. This workflow enables real-time molecular profiling, offering a transformative strategy for disease monitoring and personalised therapy in paediatric brain tumours.

## Main text

Central nervous system (CNS) tumours are the leading cause of cancer-related death in children, due to their lack of treatment options and the challenges of accurately diagnosing and monitoring disease progression^1^. Despite improvements for other paediatric cancers, mortality from childhood CNS tumours has remained largely unchanged over recent decades^2^. Conventional monitoring methods such as medical imaging and cytology are limited by false positives and low sensitivity^3-5^, which can hinder accurate disease assessment. Early and precise detection of progression or relapse is critical for timely intervention and improved outcomes.

Liquid biopsy is emerging as a promising option for CNS tumour monitoring, with current approaches focusing on cell-free DNA (cfDNA) in cerebrospinal fluid (CSF) with some studies on blood, alongside other biomarkers such as circulating RNA, extracellular vesicles and circulating tumour cells (CTCs)^6,7^. In aggressive brain tumours, tumour cells can disseminate through the CSF, leading to leptomeningeal spread, which is associated with poor prognosis and limited treatment options^8,9^. While CTCs have been studied in metastatic spread to the CSF from other cancers such as non-small cell lung cancers and breast cancer^10,11^, their role in paediatric brain tumours, particularly within the CSF, remains underexplored.

Single-cell RNA sequencing (scRNA-seq) offers an unprecedented opportunity to capture the transcriptomic landscape of individual cells, enabling molecular profiling of rare CTCs within the complex cellular environment of the CSF. By distinguishing tumour cells from normal immune, vascular, or stromal populations, scRNA-seq can reveal tumour heterogeneity, metastatic potential and therapeutic vulnerabilities. Here, we present a novel CSF processing workflow utilising scRNA-seq to sensitively detect and characterise CTCs in paediatric CNS tumours. This study lays the foundation to effectively track disease progression and potentially enable molecularly guided treatment strategies for children with brain cancer.

Sixteen patients across nine CNS tumour types enrolled on the ZERO Childhood Cancer Program (ZERO)^12^ had CSF collected as part of clinical care, with seven having multiple timepoints collected throughout their disease journey (**Fig 1A-B**). Fifteen patient’s CNS tumour biopsy had whole genome sequencing (WGS) and 13 had matched bulk RNA-sequencing from which we identified the molecular driver alterations in each tumour, including single nucleotide variants (SNV), copy number variants (CNV) and structural variants (SVs) (**Fig. 1B**).

**Figure 1.**
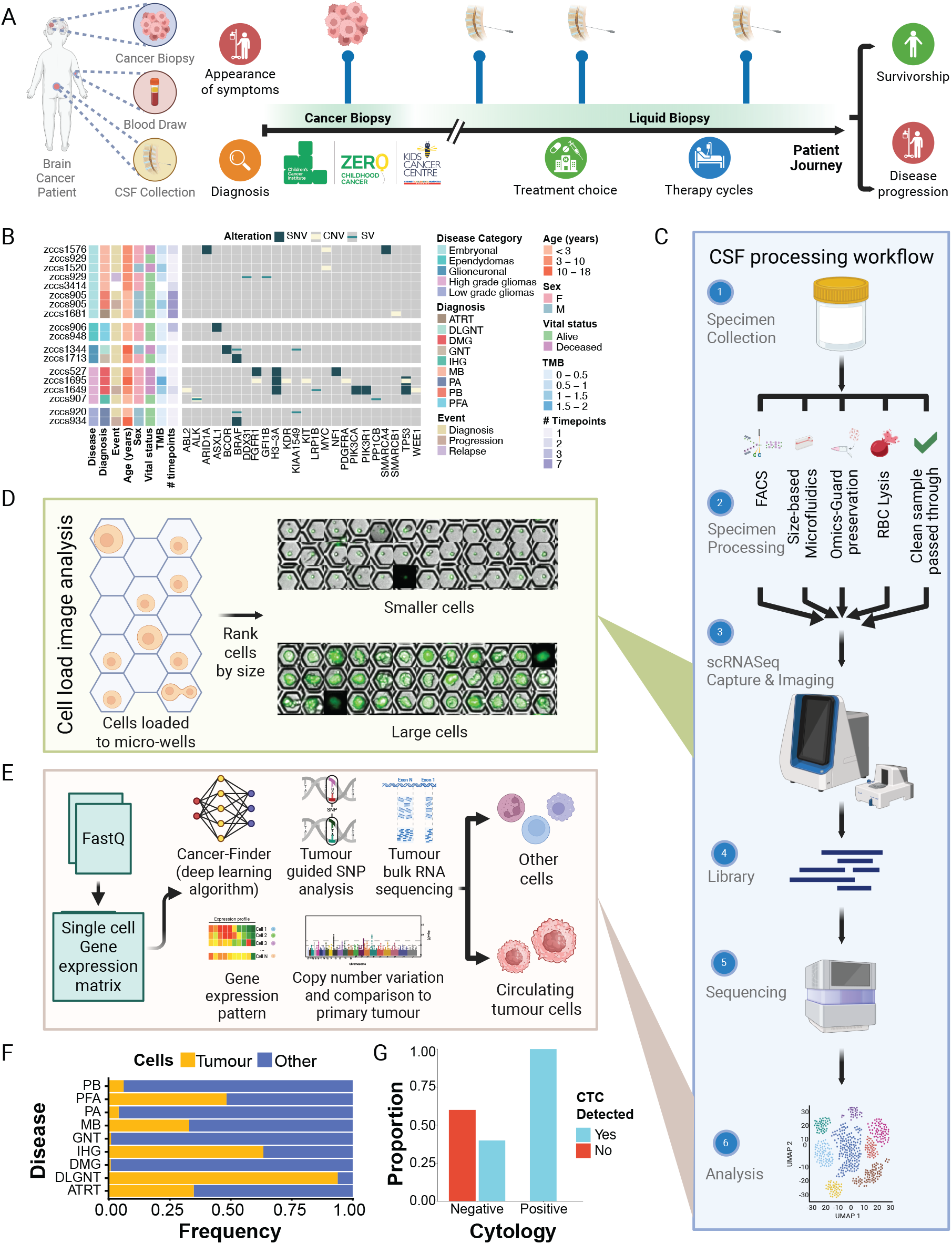
CSF processing for CTC detection using scRNA-seq in CNS tumours. **(A)** Schematic of the patient journey from diagnosis and enrolment on ZERO to sample collection, highlighting timepoints of CSF collection to aid disease progression and survivorship. **(B)** Oncoplot highlighting the mutations identified from WGS tumour samples, with patient characteristics and diagnosis annotated on the left, alongside the number of CSF samples collected. **(C)** Overview of the 6-step CSF processing workflow including: specimen collection and processing, scRNA-seq cell capture and imaging, library construction, sequencing and analysis. **(D)** Processed images generated by the BD Rhapsody™ Scanner during the Cell Load step which automates cropping and stitching of microwells containing cells and compiles them into a single image ordered by cell size. **(E)** Analytical workflow from raw fastq files to identification of non-malignant and circulating tumour cells (CTCs). Includes cancer prediction, gene expression analysis and integration of scRNA-seq with matched WGS and bulk RNA-seq from tumour molecular profiling. **(F)** Proportion of CTCs and non-malignant cells detected grouped by CNS diagnosis. **(G)** Proportion of samples that had CTCs detected from scRNA-seq analysis of the CSF in comparison to positive or negative findings as reported by standard cytology testing.

To evaluate the feasibility of scRNA-seq to identify CTCs from CSF, we first developed and optimised a dedicated CSF processing workflow. Following specimen collection, CSF samples were rapidly processed (see online methods), captured, imaged and sequenced using the BD Rhapsody™ system whole transcriptome analysis (**Fig. 1C**). Taking advantage of the integrated imaging within the BD Rhapsody™ platform, we developed an additional QC step (see online methods) that helped measure sample quality and select optimum cells by examining images of captured cells^13^, including their Calcein AM fluorescence, morphology and size (**Fig. 1D**). scRNA-seq data was analysed with cell types annotated and quantified to identify the proportion of CTCs in each sample (**Fig. 1E**). A combination of a pre-trained cancer prediction model (Cancer-Finder)^14^, tumour-informed biomarker detection of mutations (SNVs and CNVs) and marker gene expression were utilised to distinguish between CTCs and non-malignant cell clusters (**Fig. 1E**).

Our CSF processing workflow successfully detected CTCs in all patients and across different CNS diagnoses (**Fig. 1F**). Pineoblastoma (PB), medulloblastoma (MB), glial neuronal tumour (GNT), diffuse leptomeningeal glioneuronal tumour (DLGNT) and atypical teratoid rhabdoid tumours (ATRT) had the highest CTC burden, ranging from 33–94%. In contrast, posterior fossa A (PFA) ependymoma, pilocytic astrocytoma (PA), infant-type hemispheric glioma (IHG), and diffuse midline glioma (DMG) had fewer than 10% of cells annotated as CTCs (**Fig. 1F**). Importantly, our methodologies detected CTCs in 40% of samples when traditional cytology tests reported negative findings (**Fig. 1G**). Taken together, these findings confirm that our CSF workflow can sensitively detect CTCs across a spectrum of CNS tumour subtypes, even in low abundance.

Having confirmed that our methodology identifies CTCs, we next investigated whether longitudinal tracking of CTCs reflected clinical disease course. Patient zccs905 presented with signs of raised intracranial pressure including irritability, lethargy, vomiting, gait disturbance, a right abducens nerve palsy and bilateral papilloedema. Neuraxial imaging demonstrated significant hydrocephalus and a pineal mass, requiring endoscopic third ventriculostomy and tumour biopsy, which confirmed the pineoblastoma diagnosis. The patient was enrolled on ZERO where molecular profiling of the tumour at diagnosis and progression was performed (**Fig. 1B**).

To confirm accurate identification of tumour-derived CTCs and non-malignant cells, we applied both CNV prediction to annotate clusters as either aneuploid (predicted CTCs) or diploid (predicted non-malignant) and utilised Cancer-Finder^14^ to distinguish tumour and non-malignant clusters. Both identified the same clusters of CTCs and non-malignant cells (**Fig. 2A**). Orthogonal validation using immune markers (e.g. *PTPRC*) confirmed the absence of immune cells within CTC clusters, while expression of *FOXR2, MYC* and *MYCN* confirmed CTC clusters (**Fig. 2B**). These complementary strategies strengthened confidence in CTC identification.

**Figure 2.**
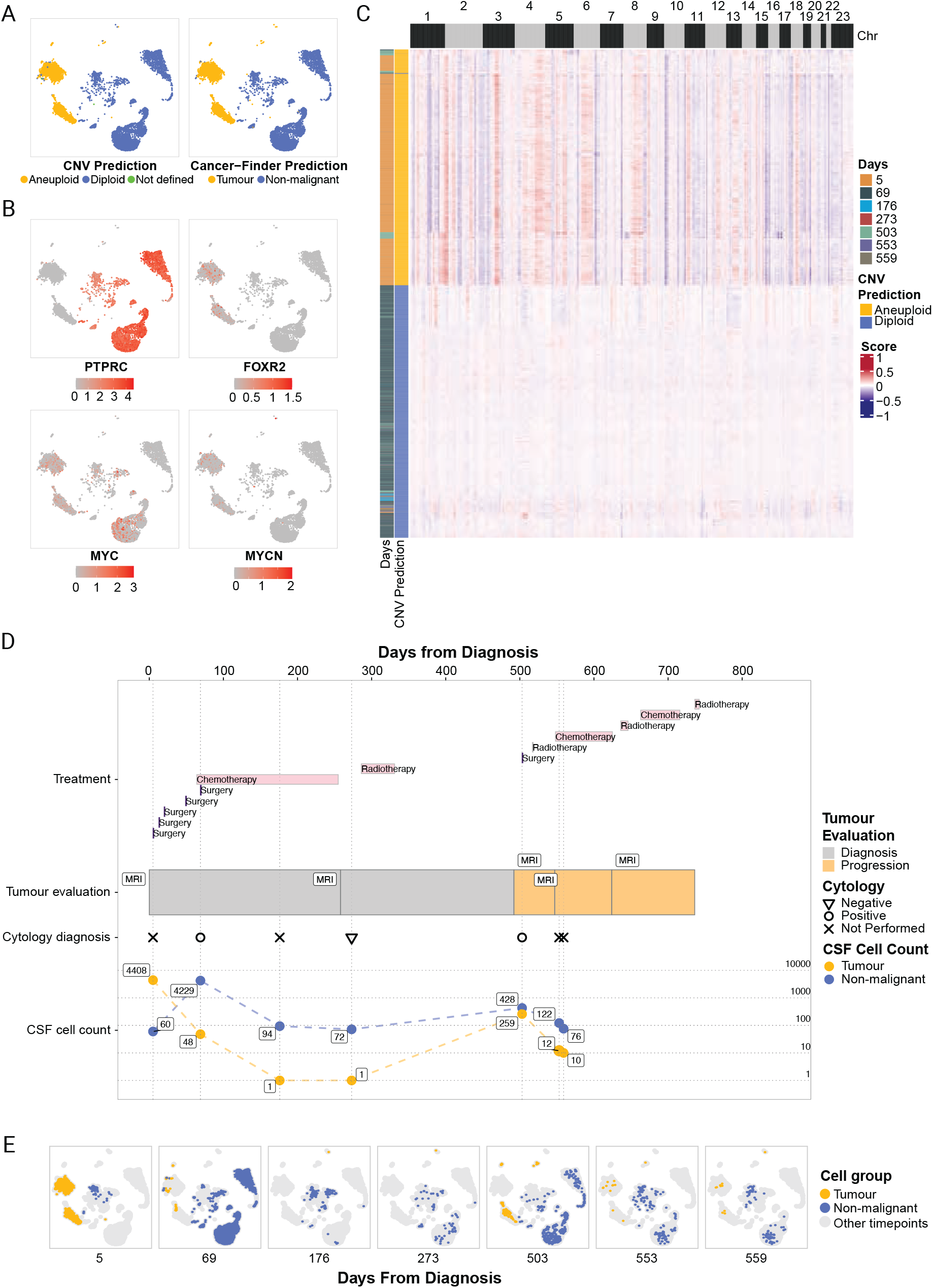
Longitudinal monitoring of a pineoblastoma patient (zccs905) Cell clustering UMAP plots highlighting **(A)** aneuploid cells indicative of CTCs (yellow) and diploid cells indicative of non-malignant cells (blue) on the left using CNV prediction and CTCs (yellow) and non-malignant cells (blue) as predicted by cancer finder on the right, and **(B)** gene expression-based markers of *PTPRC* to confirm absence of immune cells in CTC clusters, and *FOXR2, MYC* and *MYCN* markers that were elevated in tumour cells. **(C)** Copy number profile of all the cells across the seven CSF samples, separated into cells predicted to be aneuploid (CTCs) and diploid (non-malignant cells). CNV profiling revealed a sub-clonal gain of chromosome 1q at diagnosis (day 5), a hallmark of pineoblastoma, detectable only by scRNA-seq and absent in bulk RNA-seq at diagnosis. **(D)** Timeline of clinical events from top to bottom: treatment as highlighted by pink showing duration; tumour status as evaluated by MRI and interim evaluations (grey – diagnosis; yellow – progression); cytology results (triangle – negative, circle – positive and cross – not available); CSF counts of non-malignant (blue) and CTCs (red) with associated CTC counts labelled. **(E)** Cell clustering UMAP plots at matching cytology timepoints, coloured by cell type non-malignant (blue) and CTCs (red).

CSF was collected at diagnosis and serially during treatment, which included surgical debulking, chemotherapy and focal radiotherapy to the pineal tumour and one spinal nodule. Seven CSF samples were analysed using our scRNA-seq workflow, and CNV profiles enabled distinction between aneuploid (predicting CTCs) and diploid (predicted non-malignant) cells (**Fig. 2C**). At diagnosis 97% (4408/4468) of cells were detected as CTCs, consistent with high disease burden typically present at initial diagnosis (**Fig. 2D-E**). The patient underwent two debulking surgeries and extra ventricular drain (EVD) insertion which was complicated by ventriculitis. The EVD was subsequently exchanged, and a ventricular-peritoneal shunt was inserted following resolution of the ventriculitis. The patient commenced chemotherapy 69 days post diagnosis where the CSF sample was obtained and both cytology and scRNA-seq showed a subpopulation of CTCs albeit with a marked reduction. Subsequent CSF samples were collected at day 176 and 273 where cytology was negative at day 273, and scRNA-seq only revealed one CTC. The patient underwent an MRI at day 503 at which progressive disease was observed, with cytology returning a positive result and scRNA-seq demonstrated increased CTCs (**Fig. 2D–E**). CNV profiling of these CTCs revealed CNV changes consistent with those identified from WGS (chromosomes 1q gain and 8p and 16q segmental 1 copy loss). Notably, exploring the CNV changes of all serial CSF samples revealed that the diagnostic chromosome 1q gain, a hallmark of pineoblastoma, was present in a sub-clonal population of cells at initial diagnosis, that was not identified from WGS data (**Fig. 2C**). This sub-clonal CNV, which emerged using conventional approaches at progression (day 505), underscores the sensitivity of single cell approaches to identify clinically relevant alterations that would otherwise remain undetected until progression or relapse. Following additional surgery, radiotherapy and commencement of a new chemotherapy regime, two more CSF samples were collected (553 and 559 days respectively), with both samples showing a decrease in the number of CTCs present (**Fig. 2D-E**). Together, these findings demonstrate that longitudinal scRNA-seq of CSF enables precise detection of CTCs and dynamic CNV changes, providing a high-resolution view of tumour evolution and treatment response that complements conventional cytology and provides more resolution than bulk sequencing.

Our novel CSF processing and scRNA-seq workflow reliably detects and characterises CTCs, and as highlighted by following a single patient across seven CSF samples, identifies CSF resident cells as dynamic surrogates of disease state. This minimally invasive approach offers a real-time window into tumour biology, enabling refined diagnostics, monitoring of disease progression and more informed treatment decisions. Importantly, the ability to sensitively detect CTCs during routine monitoring, opens the door to earlier identification of treatment resistance or relapse, potentially before clinical symptoms emerge. This could significantly improve patient outcomes by allowing timely therapeutic intervention.

Critically, using scRNA-seq on CSF revealed a sub-clonal gain of chromosome 1q at initial diagnosis, a diagnostic marker of pineoblastoma, that bulk molecular profiling of the diagnostic tumour failed to detect. This finding underscores not only the detection of CTCs from CSF but the sensitivity of single cell methodologies to detect diagnostic clinically relevant alterations^15,16^. In contrast, a recent study implemented low-coverage WGS of CSF in paediatric brain tumours, where they identified circulating tumour DNA from CSF in only 45% of samples^17^. This highlights a key limitation of lower-resolution approaches. Our ability to identify sub-clonal events at diagnosis suggest that scRNA-seq can uncover early genomic heterogeneity with potential prognostic therapeutic implications, thus driving earlier intervention or combination therapies to reduce risk of relapse, offering a significant advance over existing genomic profiling strategies.

Beyond tumour cell identification, scRNA-seq can provide a detailed and nuanced view of the cellular makeup of the CSF throughout the disease journey. This provides insight into not only the cell types present but their activation states and potentially how they are interacting, something not possible through current standard of care testing approaches. The work presented here demonstrates the feasibility and clinical utility of longitudinal CSF sampling for CTC detection in paediatric CNS tumours. Taken together, integrating scRNA-seq based CTC detection into clinical workflows represents a transformative step in paediatric brain cancer care. It enhances our ability to monitor disease in real time, personalise treatment strategies throughout the patient’s treatment journey and ultimately improve prognosis for children with CNS tumours.

## Acknowledgements

We sincerely thank the patients and parents for participating in this study, without whom this research would not have been possible. We would also like to thank the clinicians, tumour banks and health professionals for their time acquiring consent for patients and for the collection and coordination of samples and associated clinical data at the Sydney Children’s Hospital, Randwick, and the clinical trial support staff at Koala for their vital contributions to patient care and study coordination. We thank ANZCHOG as the trial sponsor, and we thank the staff of the Clinical Translation Theme of the Children’s Cancer Institute for their dedicated work on the ZERO Childhood Cancer Program. We gratefully acknowledge the technical expertise and infrastructure provided by the Children’s Cancer Institute Technology and Innovation Enabling Platform and the Children’s Cancer Institute Bioinformatics and Research Compute Enabling Platform. The authors acknowledge the provision of computing and data resources provided by the Australian BioCommons Leadership Share (ABLeS) program. This program is co-funded by Bioplatforms Australia (enabled by NCRIS), the National Computational Infrastructure and Pawsey Supercomputing Research Centre. We thank SevenBridges for supporting the processing of raw single-cell sequencing data on their analysis platform. This work has been funded by the Australian Federal Government Department of Health, the New South Wales State Government, Cancer Institute New South Wales, Love Your Sister, the Yellow Diamond Foundation, and Tour de Cure. The Australian Cancer Research Foundation (ACRF) for funding to establish infrastructure to support the ZERO Childhood Cancer personalised medicine programme and infrastructure to support the ACRF Childhood Cancer Liquid Biopsy Program. ZERO Childhood Cancer is a joint initiative led by Children’s Cancer Institute and the Kids Cancer Centre, Sydney Children’s Hospital, Randwick. Funding from the Kids Cancer Alliance, Cancer Therapeutics Cooperative Research Centre supported the development of a personalised medicine programme; Tour de Cure supported tumour biobank personnel; the Steven Walter Children’s Cancer Foundation and the Hyundai Help 4 Kids Foundation supported P.G.E.; Samuel Nissen Charitable Foundation supported P.G.E.; The Medical Research Future Fund, Australian Brain Cancer Mission/National Health & Medical Research Council/Lifting Clinical Trials and Registry Capacity (NHMRC MRF9500002), the Minderoo Foundation’s Collaborate Against Cancer Initiative and funds raised through the ZERO Childhood Cancer Capacity Campaign, a joint initiative of Children’s Cancer Institute and the Sydney Children’s Hospital Foundation, supported the national clinical trial and associated clinical and research personnel. Cancer Institute of New South Wales and New South Wales Health (fellowship funding for M.J.C.; CINSW Program Grant 2019/TPG2037 for M.J.C and D.S.Z and 2021/TPG2112 for M.H., and L.M.S.L). National Health and Medical Research Council (Synergy Grants APP2018642 for M.H., and L.M.S.L., and APP2019056 for D.S.Z; Leadership Grant APP2017898 for D.S.Z.). The 2018 Priority-Driven Collaborative Cancer Research Scheme, co-funded by Cancer Australia and My Room, for personnel and computational support (APP1165556 awarded to M.J.C.).

## Online Methods

### Patients and Samples

Patients with central nervous system (CNS) tumours that were enrolled through the PRISM clinical trial (NCT03336931) at Sydney Children’s Hospital, Randwick, conducted as part of the Australian Zero Childhood Cancer Precision Medicine Program, that had cerebral spinal fluid (CSF) samples collected as either standard of care, during operation (if the ventricles were being accessed) or through a drain (external ventricular drain) if inserted at diagnosis were included in this study. Ethical approval was obtained through the Hunter New England Human Research Ethics Committee of the Hunter New England Local Health District (reference no. 17/02/015/4.06) and the New South Wales Human Research Ethics Committee (reference no. HREC/17/HNE/29). Informed consent for research for each participant was provided by parents/legal guardian for participants under the age of 18 years and by the participants who were over the age of 18 years. Whilst this trial was open to all eight paediatric oncology centres around Australia, samples for this study were solely collected from Sydney Children’s Hospital, Randwick, due to logistic challenges requiring processing of CSF within a four-hour window.

Molecular profiling of matched patient germline and tumour samples for whole genome sequencing (WGS) and bulk RNA-sequencing (RNA-seq) were collected and analysed as previously described^1^. For this study, we obtained the already processed single nucleotide variants (SNVs), copy number variants (CNVs), structural variants (SVs) and RNA gene expression profiles.

### CSF processing

CSF specimens collected at the Sydney Children’s Hospital, Randwick were processed within four-hours of acquisition. Red blood cell (RBC) contamination was assessed both before and after centrifugation. Depending on the degree of RBC contamination and sample arrival time, CSF samples were processed using one of four workflows: (1) samples with no visible RBCs proceeded directly to single-cell RNA-sequencing (scRNA-seq) capture; (2) samples that appeared slightly hemolyzed prior to centrifugation or exhibited a visible RBC pellet after centrifugation underwent RBC lysis using BD Pharm Lyse™ Lysing Buffer (BD Biosciences, cat. 555899); (3) samples requiring delayed processing had their cell pellets resuspended in BD OMICS-Guard Sample Preservation Buffer (BD Biosciences, cat. 570911) and were captured within 72 hours; and (4) samples with severe RBC contamination were enriched for large cells using an in-house microfluidic device or were subjected to live-cell isolation by fluorescence-activated cell sorting (FACS), selecting Calcein AM positive (BD Biosciences, cat. 564061) and DRAQ7 negative (BD Biosciences, cat. 564904) cells using a BD FACSAria™ Fusion flow cytometer.

### Cell enrichment – RBC Lysis

CSF was spun at 420g (swing-bucket rotor) for 5 minutes at 4 °C. Supernatant was removed and cells were resuspended in 500uL of 1x BD Pharm Lyse™ Lysing Buffer (BD Biosciences, cat. 555899) and incubated at room temperature for 5 minutes. Cells were centrifuged to remove the RBC lysis buffer and washed once with 2 mL of BD Pharmingen™ Stain Buffer (FBS) (BD Biosciences, cat. 554656). Cells were resuspended in the cold sample buffer from the BD Rhapsody™ Cartridge Reagent Kit (BD Biosciences, cat. 633731, 664887, or 667052) suitable for the cartridge type.

### Microfluidic device - Fabrication

The microfluidic devices were fabricated using soft lithography techniques, following standard procedures. A silicon wafer was used as the substrate for creating a master mold. First, the wafer was thoroughly cleaned using sequential rinses of acetone, isopropanol, and deionized (DI) water, followed by drying with nitrogen gas. The wafer was then subjected to a dehydration bake at 200°C for 30 minutes to remove residual moisture. Then, a SU-8 photoresist was spin-coated onto a silicon wafer to achieve the chosen height and patterned using photolithography to create a master mold with the desired zigzag channel geometry. The patterned wafer was then subjected to a post-exposure bake and developed to obtain the final mold.

For polydimethylsiloxane (PDMS) replica molding, (Sylgard 184, Dow Corning) was prepared by mixing the base polymer and curing agent at an 8:1 weight ratio. The mixture was thoroughly stirred for 10 minutes to ensure homogeneity and subsequently degassed under vacuum for 25 minutes to remove entrapped air bubbles. The degassed PDMS was then carefully poured onto the master mold, ensuring uniform coverage, and cured at 60°C for 2 hours inside an oven. Once fully cured, the PDMS layer was peeled off from the master mold, and inlet/outlet holes were punched using a biopsy punch of 1.2 mm diameter.

The PDMS replica was bonded to a clean glass slide to create enclosed microfluidic channels. To achieve irreversible bonding, both the PDMS surface and the glass substrate were treated with oxygen plasma using a plasma cleaner. Immediately after plasma activation, the PDMS was aligned and pressed onto the glass, ensuring a tight seal. The bonded device was further baked at 60°C for 2 hours to enhance adhesion.

The final devices were stored in a clean, dust-free environment until further experimentation. Also, before use, the microfluidic channels were primed by flushing with DI water to remove any residual contaminants.

### Cell enrichment – Inertial Focusing

The CSF cell suspension was loaded into a 10 mL BD plastic syringe (Becton Dickinson, Franklin Lakes, NJ, USA) and mounted onto a precision syringe pump (KD Scientific, Holliston, MA, USA). The syringe pump was programmed to deliver the fluid at controlled flow rates. The sample fluid was introduced into the microfluidic device via flexible Tygon tubing (ID: 0.02 inch, OD: 0.06 inch) to maintain a sealed flow path and minimize potential leakage. The experiments were conducted at room temperature (24°C) to ensure stable flow conditions. Microfluidic channels were visualized and recorded using an Olympus IX83 inverted epifluorescence microscope (Olympus Inc., USA). The microscope was equipped with a DP80 dual-chip CCD camera (Olympus Inc., USA) for real-time image acquisition and the PHANTOM VEO-E 340L MONO high-speed camera (Vision Research, Wayne, NJ, USA) to capture rapid particle movements. The high-speed camera was operated through Phantom Camera Control (PCC, version 3.9.40.805, 64-bit) to adjust acquisition settings such as frame rate, exposure time, and resolution. Following inertial focusing the cells within the enriched fraction (>~15microns) were collected, spun and processed for scRNA-seq.

### Cell enrichment – Flow Cytometry

Cell sorting was performed on a BD FACSAria™ Fusion configured with a 100-µm nozzle. Cells were resuspended in BD Pharmingen™ Stain Buffer (FBS) (BD Biosciences, cat. 554656) and stained with 2mM Calcein AM (BD Biosciences, cat. 564061) and 0.3mM DRAQ7 (BD Biosciences, cat. 564904) at 37 °C for 5 min in the dark, using a 1:200 dilution for each dye. Debris and non-cellular events were excluded by forward and side scatter gating. Live cells, defined as Calcein AM positive and DRAQ7 negative, were sorted in Yield mode and collected into a tube containing 500 µL of Sample Buffer from the BD Rhapsody™ Cartridge Reagent Kit.

### Cell Load image analysis

CSF samples loaded onto BD Rhapsody™ cartridges were imaged using the integrated Rhapsody Scanner module. For each capture run, the microchannel was scanned to generate 26 paired bright-field and Calcein AM (BD Biosciences, cat. 564061) fluorescence images, enabling identification of captured cells based on Calcein AM signal.

Our previously published R script^2^ was developed to extract 30 × 30-pixel regions of interest corresponding to Calcein AM positive events across the 26 image pairs. Cropped images were stratified into three size categories (0-8 μm, 8-20 μm, and >20 μm) and, within each category, bright-field and fluorescence crops were stitched into composite images with cells ordered by increasing size. For each run, the script generated a data frame containing cell size measurements, spatial coordinates within the cartridge, and singlet versus multiplet annotations. These features were subsequently used to derive cell-size distributions and violin plots for quality assessment. More information and the script for this analysis were described previously^2^.

### scRNA-seq processing and analysis

CSF-derived cells were processed and captured using the BD Rhapsody™ platform with the BD Rhapsody™ Cartridge Kit (BD Biosciences, cat. 633733 or 666262), the BD Rhapsody™ Cartridge Reagent Kit (BD Biosciences, cat. 633731, 664887, or 667052), and the BD Rhapsody™ cDNA Kit (BD Biosciences, cat. 633773). Sequencing libraries were prepared using the BD Rhapsody™ Whole Transcriptome Analysis (WTA) Amplification Kit (BD Biosciences, cat. 633801). Libraries were sequenced on a NovaSeq 6000 or NovaSeq X platform (Illumina) at the Ramaciotti Centre for Genomics, University of New South Wales, with a target sequencing depth of 60,000 reads per cell.

Sequencing reads from each CSF sample were processed using the BD Rhapsody™ Sequencing Analysis Pipeline (v2), implemented on the Seven Bridges cloud-computing platform (www.sevenbridges.com). In addition to the raw fastq files, the BD Bioinformatics reference package RhapRef_Human_WTA_2023-03.tar.gz was provided for alignment of human samples. Default pipeline parameters were used, including alignment to both exonic and intronic regions, with the exception that generation of corresponding BAM files was enabled. Gene expression matrices produced by the pipeline were converted into Seurat^3-7^ objects for downstream analysis. Cells with fewer than 200 detected genes were removed prior to downstream analysis. For each cell, the percentages of mitochondrial and haemoglobin gene expression were calculated. Given the stringent Cell Lysis step in the BD Rhapsody™ workflow, cells exhibiting more than 40% mitochondrial gene expression were excluded. Putative red blood cells, defined by haemoglobin gene expression exceeding 1%, were also removed.

Following these initial filtering steps, each gene expression matrix underwent three rounds of automated cell annotation: doublet detection, cell type prediction, and malignancy assessment. Cell type annotation was first performed using SingleR^8^ with Celldex reference datasets, specifically the Blueprint Encode and Human Primary Cell Atlas collections. Additional cell type predictions were generated using Azimuth^4^ with the PBMC reference and CellTypist^9^ with the pre-trained Immune_All_Low.pkl and Developing_Human_Brain.pkl models. Cells expressing *PPBP* (>0 counts) that were concurrently classified as platelets by SingleR (Human Primary Cell Atlas reference) and Azimuth were removed. Doublets were identified using scDblFinder^10^ and excluded. For malignancy prediction, each count matrix with gene symbols as row names and cell identifiers as columns was processed with Cancer-Finder^11^ and the pre-trained model model_epoch92.pkl. The resulting malignancy predictions were incorporated into the corresponding Seurat metadata. The 33 Seurat objects were merged into a single Seurat object and processed using the standard Seurat workflow with default parameters. This included data normalization (NormalizeData), identification of highly variable features (FindVariableFeatures), scaling and centring of selected features, and principal component analysis on the scaled data.

Batch-effect correction was performed using IntegratedLayers with the HarmonyIntegration method, which generated a dimensionality reduction embedding labelled “harmony.” This harmony reduction was subsequently used in FindNeighbors to compute nearest-neighbour relationships and construct the shared nearest-neighbour graph. Clustering was then carried out using FindClusters. For visualization, uniform manifold approximation and projection (UMAP) was computed on the integrated dataset.

scRNA-seq count matrices from the merged datasets were subsetted to extract cells only from patient zccs905. To infer CNV profiles the count matrix was analysed using the copykat R package. Key parameters were set as follows: id.type = “S”, cell.line = “no”, ngene.chr = 5, win.size = 25, KS.cut = 0.1, distance = “euclidean”, and genome = “hg20”. CNV heatmap was generated using the ComplexHeatmap package with the relative copy number signals as input with hierarchical clustering used for row ordering.

